# Cultural and morphological divergence of Darwin’s cactus finches (*Geospiza scandens*) across Galápagos Islands

**DOI:** 10.1101/2025.06.18.660308

**Authors:** Melanie Kaluppa, Jefferson García-Loor, Alper Yelimlieş, Çağlar Akçay, Sonia Kleindorfer

## Abstract

Understanding the divergence of cultural traits, such as bird song, provides critical insights into evolutionary processes. In geographically isolated populations, the divergence of traits associated with mate choice can lead to further genetic separation. This study investigates divergence in song syllable types, acoustic characteristics, singing behavior, and morphology in two allopatric populations of Darwin’s cactus finch (*Geospiza scandens*) on Floreana and Santa Cruz Islands in the Galápagos Archipelago. Using song recordings of 50 males, we identified 25 syllable types with no overlap between islands, indicating a complete divergence in syllable repertoires. Syllables of Floreana and Santa Cruz males diverged in the acoustic space largely due to broader frequency bandwidths on Floreana. Moreover, Floreana males had smaller beaks than Santa Cruz males. Despite acoustic and morphological divergence, singing behaviors - like syllable repetition rate and number of syllables per song - did not differ significantly. Both cultural processes, including drift and transmission biases, as well as selection on morphology could have contributed to the observed acoustic divergence. This study adds to a growing literature on the role of geographic separation in the accumulation of both cultural and morphological divergence between populations. Future research could interrogate speciation scenarios in cactus finches if populations cease to interbreed after secondary contact.

## Introduction

Allopatric speciation, the process by which geographically isolated populations diverge into distinct species, is a cornerstone of evolutionary biology (Mayr, 1942). Physical barriers restrict gene flow between populations, which can lead to independent evolutionary pathways and the accumulation of morphological and behavioral differences. When such differences influence mate choice, they can contribute to reproductive isolation, and, ultimately, speciation (Mayr, 1963; Coyne & Orr, 2004). Among behavioral traits, song in songbirds has been widely studied as a mechanism for maintaining barriers to gene flow. Birdsong plays a central role in both species recognition and mate choice (Catchpole & Slater, 2008; Price, 2008) and it reflects both cultural and genetic evolution, shaped by social learning and morphological constraints (Nowicki & Searcy, 2005; Searcy *et al*., 2021; Vernes *et al*., 2021).

Bird song learning involves the cultural transmission of songs through a process of exposure, attention, imitation, and reproduction (Catchpole & Slater, 2008). These processes offer numerous opportunities for error or modification of songs. The individual learning processes that lead to these changes can interact with social learning biases which would lead to cultural evolution (Lipkind & Tchernichovski, 2011; Aplin, 2019; Williams & Lachlan, 2022). It is worth noting that cultural evolution—defined as the transmission of behavioral traits across generations through social learning (Klump *et al*., 2021)—is documented in vocal production learning systems beyond songbirds (oscine passerines), including at least two other orders of birds (parrots (Smith-Vidaurre, Perez-Marrufo, & Wright, 2021) and hummingbirds (González & Ornelas, 2014)), cetaceans (Garland & McGregor, 2020), bats (Lin *et al*., 2015), and seals (Reichmuth & Casey, 2014). Among these, bird song remains the most prominent models for cultural evolution in vocalizations (Whiten, 2019).

Since the first systematic study of the songs of common chaffinch (*Fringilla coelebs*) by Marler (1952), geographic variation in birdsong has been documented across a wide range of species and spatial scales (Galeotti, Appleby, & Redpath, 1996; Kroodsma, 1985; Leader, Wright, & Yom-Tov, 2000; Slabbekoorn, Jesse, & Bell, 2003; Reyes et al., 2024). Such variation typically results from a combination of geographic isolation and the dynamics of song learning (Podos & Warren, 2007). Over time, cultural evolution can drive the emergence of new song types and the formation of local dialects, resulting in population-specific vocal patterns (Slater, 2003; Parker, Hauber, & Brunton, 2010; Aplin, 2016). For example, isolated populations of swamp sparrows (*Melospiza georgiana*) maintain distinct song variants through cultural transmission, even in the absence of environmental or ecological differences (Lachlan et al., 2018). These vocal differences can act as prezygotic barriers by reducing interbreeding between populations, demonstrating how cultural evolution may contribute to reproductive isolation and, ultimately, speciation (Grant & Grant, 1996; Colombelli-Négrel & Kleindorfer, 2021).

In addition to the processes that underpin cultural evolution, in vocal-learning birds, morphological traits, such as those associated with beak shape, impose biomechanical constraints on sound production, creating a link between morphology and song structure (Podos & Nowicki, 2004; Huber & Podos, 2006; Christensen, Kleindorfer, & Robertson, 2006; Podos, Lahti, & Moseley, 2009). Birds with larger beaks often produce slower or less complex songs, likely due to biomechanical constraints that make it more difficult to filter out harmonics in rapid or intricate vocalizations (Podos, 2001). During singing, birds modify the length of their vocal tract - and consequently its resonant frequencies - by opening and closing their beaks to emphasize different frequencies (Riede *et al*., 2006). Larger beaks, however, may be slower to open and close, limiting the speed and precision with which frequency modulation can occur. As a result, birds with larger beaks are often constrained to produce songs with narrower frequency bandwidths and slower repetition rates (Podos, 2001; Podos, Southall, & Rossi-Santos, 2004; Huber & Podos, 2006). Additionally, body size can also influence song characteristics (Ryan & Brenowitz, 1985; Grant, 1986; Brumm, 2009; Martin *et al*., 2011). Body size affects syrinx volume, which in turn influences the minimum fundamental frequency of the song, as demonstrated in Darwin’s finches (Bowman, 1983) and other bird species (Badyaev & Leaf, 1997; Kirschel, Blumstein, & Smith, 2009).

Darwin’s finches of the Galápagos Islands represent a particularly compelling model for studying the dynamics of speciation. Finch song typically consists of repeated renditions of one to two different syllable types, which are culturally transmitted. Juvenile males learn these syllables from a male tutor, usually their father or a nearby conspecific (Bowman, 1979, 1983; Millington & Price, 1985; Grant & Grant, 1996, 2018; Christensen & Kleindorfer, 2007; Peters & Kleindorfer, 2018). Once learned, a finch’s syllable type(s) generally remain unchanged throughout its adult life, as observed in ground finches (Gibbs, 1990; Grant & Grant, 1996) and in male small tree finches (*Camarhynchus parvulus*) over a two-year period (Christensen *et al*., 2006). Darwin’s finches exhibit song divergence across geographically separated populations (Grant & Grant, 1997a, 1997b; Colombelli-Négrel & Kleindorfer, 2021; Colombelli-Négrel et al., 2023), and this divergence has been linked to evolutionary and speciation processes (Grant & Grant, 2009; Podos & Schroeder, 2024). Beyond song divergence, Darwin’s finches show substantial morphological variation in beak and body size, not only between species but also between populations (Grant *et al*., 1985) and these correlate with vocal signal structure like frequency bandwidth and trill rate (Bowman, 1979; Podos, 2001). The combination of vocal variation and morphological diversity makes Darwin’s finches an excellent system for exploring the interplay between cultural drift, morphology, and acoustic divergence.

In this study, we investigate potential differences in song by identifying the range of song syllable types and differences in acoustic characteristics of syllables, male singing behavior, and male morphology across two geographically separated populations of Darwin’s cactus finches (*Geospiza scandens*) on Floreana and Santa Cruz Islands in the Galápagos Archipelago. First, we compile a library of syllable types within each population. Second, we compare the acoustic characteristics of syllable types between the two island populations. Third, we analyze differences in male singing behavior and, fourth, we assess morphological measurements across populations. If cultural divergence in song exists between the two islands, we predict 1) both shared and unshared (population-specific) syllable types on each island, and 2) inter-island differences in the acoustic characteristics of syllable types. Additionally, if there are differences in song production or morphological constraints influencing song, we predict: (3) inter-island differences in singing behavior—specifically in the number of syllable types per song, the number of syllables per song, and syllable repetition rate—and (4) inter-island differences in morphological measurements.

## Material & Methods

### Study sites & species

We collected acoustic data during the breeding season in March 2024, with recording sites located in the lowlands of Santa Cruz (Puerto Ayora: 0°44’S, 90°18’W, Garrapatero: 0°40’S, 90°13’W) and Floreana Island (1°,16’S, 90°29’W) in the Galápagos Archipelago (Figure 1). The morphological data were collected between 2000 and 2024 as part of a long-term mist-netting (Supporting Information, Table S1). Located in the centre of the archipelago, Santa Cruz is the second largest island (986km²) and lies approximately 47 km from the smaller Floreana Island (∼173 km²). The arid lowland sites on both islands are primarily characterized by vegetation including shrubs, dwarf trees, and cacti from the genera *Opuntia* and *Jasminocereus* (Candelabra cactus) (Jackson, 1993).

**Figure 1.**
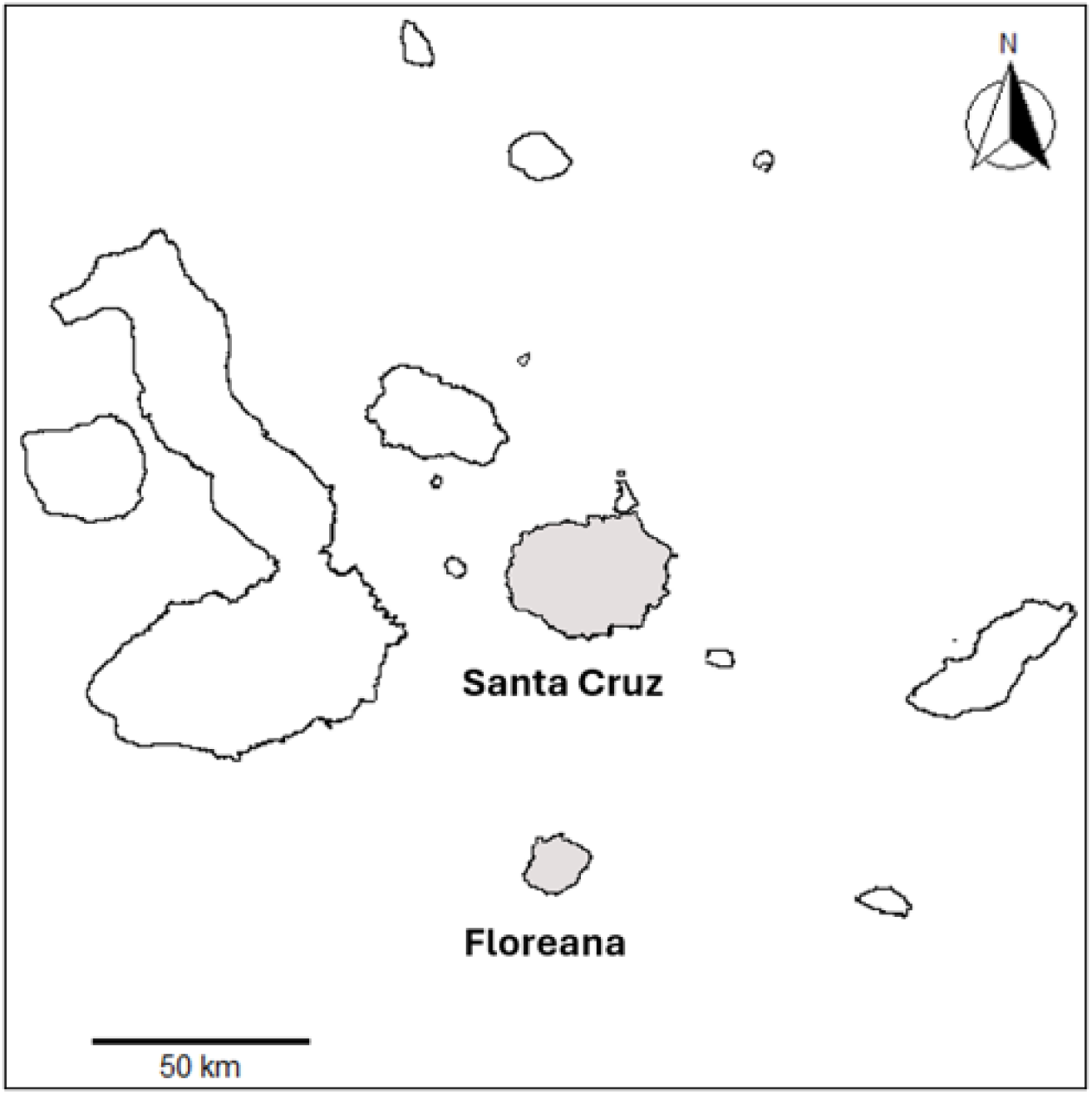
Map of the Galápagos Archipelago, with Santa Cruz and Floreana highlighted. The two islands are located approximately 47 km apart.

The common cactus finch (*Geospiza scandens*), one of 17 Darwin’s finch species (Passeriformes: Thraupidae), is a medium-sized ground finch (∼22g) found on most major Galápagos Islands, including Pinta, Santa Fé, Floreana, Santa Cruz, Isabela, Pinzón, Marchena, Santiago, and Rábida. Where absent, it is typically replaced by its close relatives — on Española, for example, by the Española cactus finch (Kleindorfer *et al*., 2019a; Jaramillo & D. A. Christie, 2020). The common cactus finch inhabits dry scrublands and arid lowland woodlands, closely associated with prickly pear cacti (*Opuntia*) for food and nesting (Boag & Grant, 1984). Its long beak, which is deep at the base and tapers to a point, is adapted for feeding on *Opuntia* pulp, seeds, and flowers (Abbott, Abbott, & Grant, 1977; Grant & Grant, 1981; Millington & Grant, 1983). Breeding typically begins in January or February, following the onset of heavier rains (Boag & Grant, 1984). During this period, males build doom-shaped display nests and sing actively to attract mates and defend territories (B. R. Grant & Grant, 1996).

### Recordings

To examine syllable type occurrence, song characteristics, and singing behavior between the two cactus finch populations, we recorded 25 males on each island, totaling 50 individuals. For 14 of the 25 males on Floreana, and all 25 males on Santa Cruz, we used playback of sympatric conspecific song to elicit singing (for the target male’s song recording) when males were observed within their territory but were not spontaneously vocalizing. Playback consisted of a maximum of six songs per trial. The one-minute playback stimuli to elicit singing were broadcast with a JBL Clip 4 Bluetooth speaker (JBL Inc., USA) and consisted of six evenly spaced songs from the same individual, high-pass filtered at 1500 Hz, with normalized peak amplitude, and saved as 16-bit WAV files. The stimulus songs were recorded in the days prior, while the first individuals were establishing their territories. We accounted for playback use in our analyses to control for potential influences on measured song parameters. Recordings were made using a Zoom F3 Field Recorder (Zoom Corporation, Japan) and a Sennheiser MKE 600 directional microphone (Sennheiser electronic GmbH & Co. KG, Germany) at a sampling rate of 48 kHz and 32-bit float resolution. We aimed to record at least 10 high-quality songs per male (Floreana: mean = 11.7, SD = 2.94; Santa Cruz: mean = 14, SD = 3.20).

### Acoustic analysis

All audio recordings were initially visualized in Audacity 3.4.2 (Audacity Team, 2024) and exported at a 48 kHz sampling rate and 16-bit depth. Spectrograms were created using Raven Pro 1.6.5 (K. Lisa Yang Center for Conservation Bioacoustics at the Cornell Lab of Ornithology, 2024) with the following settings: Hann window, window length = 512, contrast = 50, brightness = 50. For analysis, we selected high signal-to-noise ratio songs, yielding 642 total songs (Floreana: 292, Santa Cruz: 350). To define the start and end of each song, we classified a vocalization as a song if the time interval between the last syllable of the previous song and the first syllable of the next song was minimum one second apart. Doubling this threshold from one to two seconds affected only 7 out of 635 songs.

Each song was saved as an individual file and analyzed in RStudio (R Core Team, 2024). We used the *warbleR* (Araya-Salas & Smith-Vidaurre, 2017) and *ohun* (Araya-Salas & Smith-Vidaurre, 2017) packages to apply energy-based detection through the *energy_detector()* function, which marked syllable start and end points within each song, based on energy and temporal features. To prevent low-frequency noise from affecting relative amplitude, a bandpass filter (1.5–8 kHz) was applied during detection. Detected syllables were manually verified on labeled spectrograms using *label_spectro()*, adjusting *threshold*, *smooth*, and *hold.time* as needed. This process identified a total of 2001 syllables (Floreana: 888, Santa Cruz: 1113).

To extract frequency- and time-based parameters for each syllable, we first used *freq_ts()* to obtain the dominant frequency contour as a time series, from which minimum and maximum dominant frequencies, bandwidth, and slope were derived. Additional features were calculated via *spectro_analysis()*, including duration, mean/median frequency, standard deviation, first/third quartile frequencies and times, interquartile ranges (frequency/time), skewness, kurtosis, and spectral, temporal, and spectrographic entropy (see package manual for details). Syllables were filtered (>1500 Hz), and peak frequency was derived using *meanspec()* from the *seewave* package (Sueur, Aubin, & Simonis, 2008), representing the frequency of highest amplitude of each sound file. Song duration was calculated from the start of the first to the end of the last detected syllable during energy-based syllable detection. Syllable repetition rate was computed by dividing the number of syllables per song by the song duration.

### Categorization of syllable types

Using the spectrograms generated in Raven Pro 1.6.5, two researchers first categorized the syllables into syllable types by consensus on the basis of structural similarities. To assess the generalizability of this categorization across researchers, five other raters were asked to classify the syllable types into categories. For this task, we used R’s *spectro()* function from the *seewave* package to create spectrograms at the same scale, enhancing background-to-signal contrast. The raters received two printed spectrograms per syllable type, accompanied by instructions to create unique syllable type categories and/or match syllable type pairs for each island based on perceived image similarity. Inter-rater agreement was assessed by calculating the proportion of correctly matched syllable types between each rater and the original classification. The average proportional agreement across all raters was then calculated to provide an overall measure of inter-rater reliability.

### Morphological measurements

To examine morphological differences between the islands, we analyzed measurements from 36 cactus finch males (17 from Floreana and 19 from Santa Cruz) at the time of capture and banding as part of our long-term mist-netting between 2000 and 2024 (Supporting Information, Table S1). The measured parameters included beak length culmen (tip of beak to base of feathers, mm), beak width measured at the culmen (mm), beak depth measured at the culmen (mm), tarsus length (mm), and flattened wing length (mm).

## Data analysis

### Acoustic structure

We conducted a principal component analysis (PCA) using 21 frequency- and time-related measurements with *prcomp()* from R’s base *stats* package (R Core Team, 2024).The first four principal components (PC1–PC4), each with an Eigenvalue greater than 1, collectively explained 86% of the variance and were selected for further analysis (Supporting Information, Table S2). PC factor loadings for all 21 acoustic variables were calculated (Supporting Information, Table S3).

PC1–PC4 were then used to conduct linear mixed models (LMMs), with each principal component as the response variable using *lmer()* from the package *lme4* (Bates *et al*., 2015). Island and playback use (yes/no) were included as fixed effects, while bird ID and syllable type were included as random effects.

### Singing behavior

To evaluate potential differences in male singing behavior between the two islands, we analyzed the number of syllable types per song, the number of syllables per song and the syllable repetition rate.

Generalized linear mixed models (GLMMs) from the *lme4* and *glmmTMB* packages (Bates *et al*., 2015, Brooks *et al*., 2017)) were used for the number of syllable types per song and the number of syllables per song as response variables, with island and playback use included as fixed effects. Random effects differed by model: for the number of syllables per song, both bird ID and syllable type were included, while for the number of syllable types per song, only syllable type was used due to negligible variance attributable to bird ID. A generalized Poisson error distribution was applied to the GLMM of the number of syllables to account for underdispersion. We modeled the number of syllable types as a binary response because our data contained only ones and twos. We subtracted one from each row and used a binomial response in our model. To assess variations in the syllable repetition rate between island populations, a LMM (package *lme4*) was applied, including island and playback as fixed effects and bird ID and syllable type as random effects.

### Morphology

A principal component analysis (PCA) was conducted on three beak measurements (length, width, and depth). The first principal component (PC1) had an eigenvalue greater than 1 and explained approximately 63% of the total variance; it was therefore used as the primary composite measure of beak size (Supporting Information, Table S4). A separate PCA on body measurements (tarsus and wing length, in mm) identified PC1 as the main body size metric, explaining 62% of the variance (eigenvalue = 1.23; Supporting Information, Table S4). Both PC1_beak and PC1_body were normally distributed and exhibited homogeneity of variance, as confirmed by the Shapiro– Wilk and Levene’s tests, respectively. We then performed independent two-tailed *t*-tests on PC1_beak and PC1_body to evaluate morphological differences in males between the two island populations.

## Results

### Syllable types

Visual inspection of spectrograms identified 12 syllable types on Floreana and 13 syllable types on Santa Cruz, yielding a total of 25 syllable types. Classification was validated by five independent raters, resulting in an overall inter-rater agreement of 97%. Syllable types 1–12 were unique to Floreana Island, while types 13–25 were exclusive to Santa Cruz Island. No syllable types were shared between the two island populations (Figure 2).

**Figure 2.**
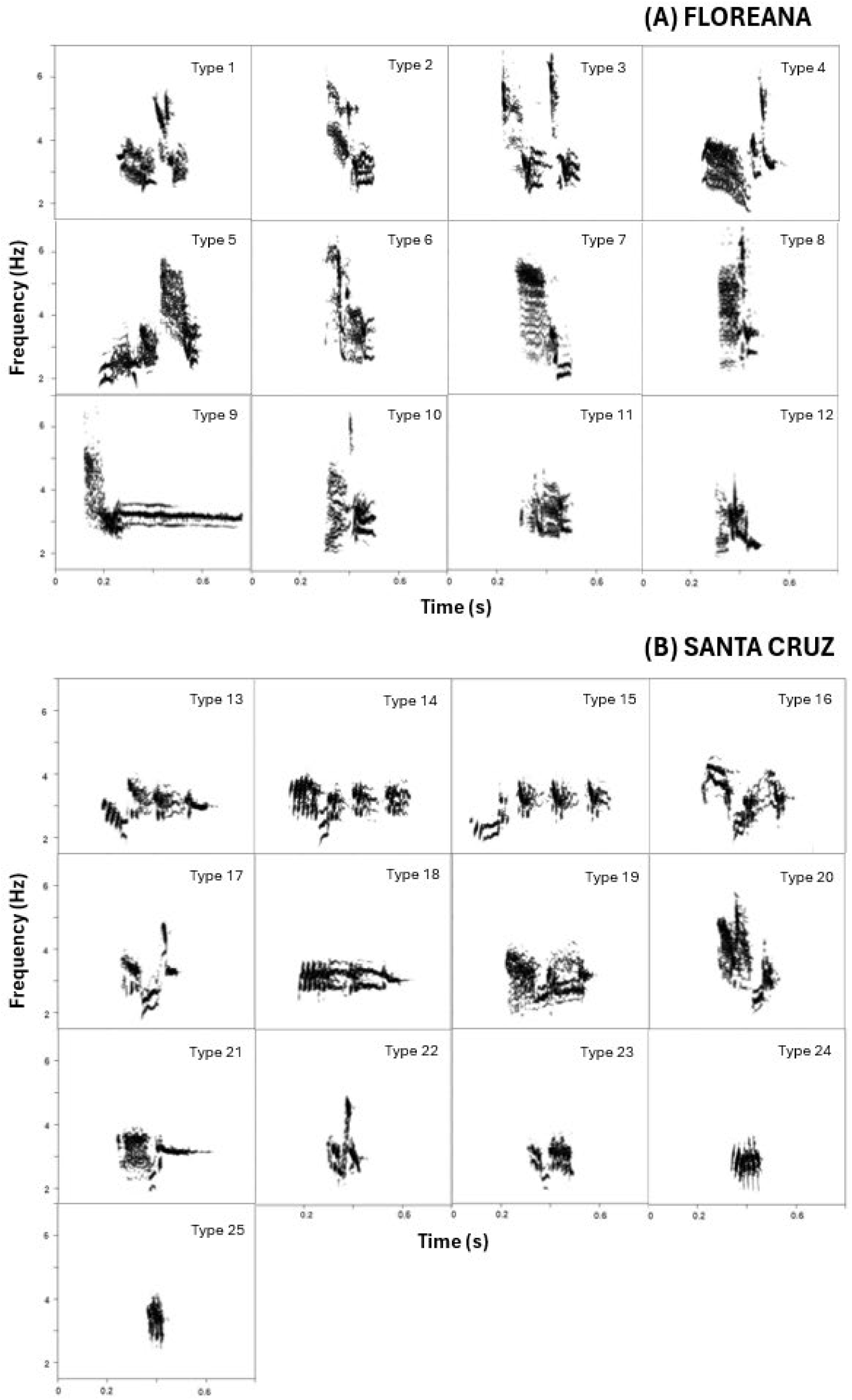
Examples of syllable types on **(A) Floreana** and **(B) Santa Cruz**: Syllable types 1–12 were found exclusively on Floreana Island, while syllable types 13–25 were unique to Santa Cruz. No syllable types were shared between the islands.

Of the 50 recorded males, 7 individuals (14%) produced songs containing two distinct syllable types, occurring in 12–100% of their songs. The remaining 43 males (86%) produced songs consisting of only one syllable type. Figure S1 (Supporting Information) presents spectrogram examples of songs from males with two syllable types. On Floreana Island, 5 out of 25 males exhibited songs with two syllable types, compared to 2 out of 25 males on Santa Cruz Island (Supporting Information, Table S5).

On Floreana Island, syllable type 2 occurred in the same song as syllable type 11 in two males, while three other males sang only syllable type 2 in their songs. On Santa Cruz, syllable type 13 occurred in the same song as syllable type 24 in one male and was also used by another male who sang only that syllable type. The syllable types 14 and 15 were produced separately in different songs by two males each, and both types occurred within the same song in one male.

On Floreana Island, half of the syllable types (6 out of 12) were each produced by 2–5 males, while the remaining six syllable types were each sung by only one male (Table 1). On Santa Cruz Island, over half of the syllable types (8 out of 13) were each produced by 2–6 males, with the remaining five syllable types each sung by a single male (Table 1).

**Table 1.** Characteristics of syllable types on **(A) Floreana** and **(B) Santa Cruz** Islands. Mean values and standard deviation of the measured parameters including duration, minimum dominant frequency, maximum dominant frequency, bandwidth and peak frequency. Data include 25 males per island whereby 5 males had songs with two syllable types on Floreana Island and two males had songs with two syllable types on Santa Cruz. Syllable types are listed in order of their percentage occurrence, with the number of males producing each syllable type indicated in parentheses.

**(A) FLOR EANA**
| Syllable type | % occurrence | Duration (s)<sup>a</sup> | Minimum dominant frequency (kHz)<sup>a</sup> | Maximum dominant frequency (kHz)<sup>a</sup> | Bandwidth (kHz)<sup>a</sup> | Peak frequency (kHz)<sup>a</sup> |
| --- | --- | --- | --- | --- | --- | --- |
| 1 | 16.7 (5) | 0.28 (0.02) | 2.61 (0.12) | 5.49 (0.61) | 2.88 (0.69) | 3.13 (0.19) |
| 2 | 16.7 (5) | 0.22 (0.04) | 2.66 (0.08) | 6.02 (0.53) | 3.36 (0.53) | 3.52 (0.89) |
| 12 | 16.7 (5) | 0.21 (0.03) | 2.25 (0.08) | 4.3 (0.53) | 2.05 (0.53) | 2.68 (0.56) |
| 11 | 13.3 (4) | 0.14 (0.03) | 2.72 (0.14) | 3.79 (0.30) | 1.07 (0.28) | 3.1 (0.29) |
| 3 | 10.0 (3) | 0.31 (0.02) | 2.7 (0.12) | 6.18 (0.28) | 3.48 (0.32) | 3.3 (0.09) |
| 5 | 6.7 (2) | 0.42 (0.01) | 1.99 (0.18) | 5.32 (0.15) | 3.33 (0.25) | 2.54 (0.15) |
| 4 | 3.3 (1) | 0.33 (0.02) | 2.08 (0.13) | 5.69 (0.25) | 3.61 (0.31) | 3.38 (0.22) |
| 6 | 3.3 (1) | 0.21 (0.01) | 2.73 (0.03) | 6.09 (0.18) | 3.36 (0.19) | 3.51 (0.14) |
| 7 | 3.3 (1) | 0.21 (0.01) | 2.15 (0.04) | 5.33 (0.10) | 3.18 (0.12) | 4.69 (0.75) |
| 8 | 3.3 (1) | 0.17 (0.01) | 2.98 (0.30) | 6.04 (0.68) | 3.06 (0.55) | 3.53 (0.36) |
| 9 | 3.3 (1) | 0.6 (0.06) | 2.78 (0.13) | 5.05 (0.27) | 2.27 (0.31) | 3.25 (0.06) |
| 10 | 3.3 (1) | 0.2 (0.01) | 2.57 (0.12) | 5.3 (0.72) | 2.73 (0.74) | 2.84 (0.23) |
| 100.0 (30) |  |  |  |  |  | <sup>a</sup>Mean (SD) |

**(B) SANT A CRUZ**
| Syllable type | % occurrence | Duration (s)<sup>a</sup> | Minimum dominant frequency (kHz)<sup>a</sup> | Maximum dominant frequency (kHz)<sup>a</sup> | Bandwidth (kHz)<sup>a</sup> | Peak frequency (kHz)<sup>a</sup> |
| --- | --- | --- | --- | --- | --- | --- |
| 22 | 22.2 (6) | 0.16 (0.03) | 2.59 (0.26) | 4.09 (0.65) | 1.5 (0.75) | 3.2 (0.18) |
| 16 | 11.1 (3) | 0.33 (0.03) | 2.26 (0.18) | 4.28 (0.15) | 2.02 (0.18) | 3.67 (0.50) |
| 23 | 11.1 (3) | 0.21 (0.02) | 2.31 (0.11) | 3.33 (0.12) | 1.02 (0.15) | 3.09 (0.09) |
| 13 | 7.4 (2) | 0.45 (0.03) | 2.38 (0.12) | 3.61 (0.30) | 1.23 (0.25) | 3.04 (0.08) |
| 14 | 7.4 (2) | 0.48 (0.02) | 2.36 (0.09) | 3.68 (0.06) | 1.33 (0.11) | 3.18 (0.16) |
| 15 | 7.4 (2) | 0.51 (0.05) | 2.1 (0.16) | 3.57 (0.16) | 1.47 (0.25) | 3.18 (0.12) |
| 17 | 7.4 (2) | 0.27 (0.02) | 2.28 (0.16) | 4.72 (0.30) | 2.44 (0.32) | 3.33 (0.17) |
| 18 | 7.4 (2) | 0.45 (0.04) | 2.47 (0.29) | 3.34 (0.07) | 0.87 (0.24) | 3.04 (0.14) |
| 19 | 3.7 (1) | 0.35 (0.01) | 2.47 (0.05) | 3.76 (0.12) | 1.29 (0.13) | 3.21 (0.20) |
| 20 | 3.7 (1) | 0.24 (0.01) | 2.29 (0.09) | 4.93 (0.10) | 2.64 (0.10) | 3.56 (0.52) |
| 21 | 3.7 (1) | 0.28 (0.02) | 2.4 (0.11) | 3.59 (0.12) | 1.19 (0.17) | 3.13 (0.03) |
| 24 | 3.7 (1) | 0.11 (0.01) | 2.94 (0.23) | 3.68 (0.33) | 0.74 (0.25) | 3.51 (0.32) |
| 25 | 3.7 (1) | 0.08 (0.01) | 3.09 (0.17) | 3.65 (0.13) | 0.56 (0.19) | 3.3 (0.08) |
| 100.0 (27) |  |  |  |  |  | <sup>1</sup>Mean (SD) |

### Differences in the acoustic structure of the syllable types across islands

The linear mixed models showed significant differences between islands for PC1 (higher factor loadings for frequency bandwidth) and PC4 (dominated by peak frequency). However, after applying a false discovery rate (FDR) correction (Benjamini-Hochberg) to account for multiple comparisons across PC1–PC4, the inter-island difference in PC4 was no longer statistically significant (adjusted p = 0.113). A robust and significant difference remained for PC1. Specifically, males from Floreana produced syllable types with broader frequency bandwidths, due to more energy in higher frequencies, compared to males from Santa Cruz (PC1 difference estimate = 1.78, SE = 0.40, t = 4.42, p < 0.001). No significant differences were found between the islands for PC2 (time-related variables) or PC3 (spectral shape features). Playback had no significant effect on the acoustic features measured.

Visualization of PC1 and PC2 revealed not only differences between islands (Figure 3, A) but also variation among syllable types overall. Each syllable type clustered recognizably according to PC1 (frequency bandwidth) and PC2 (time) (Figure 3, B).

**Figure 3.**
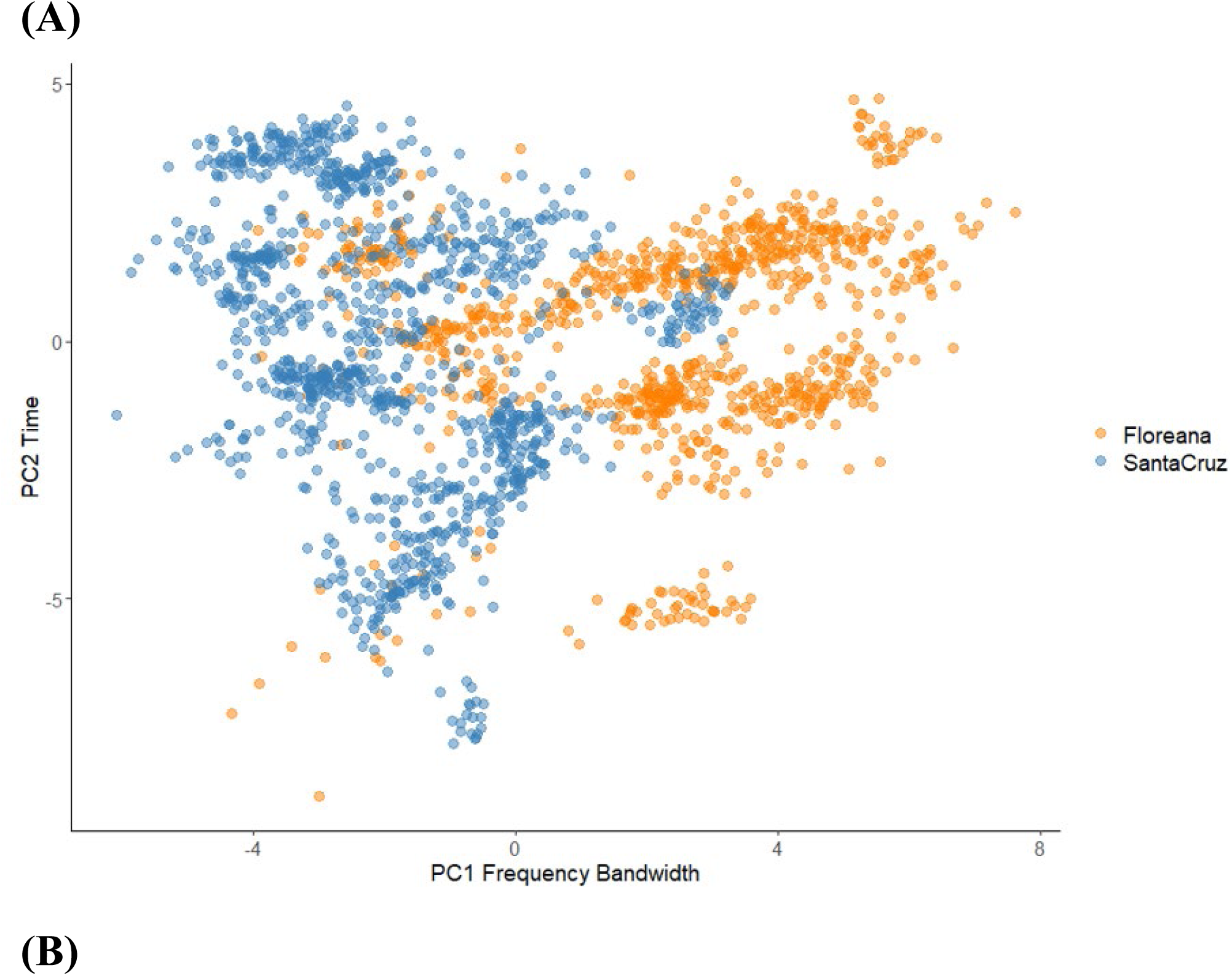
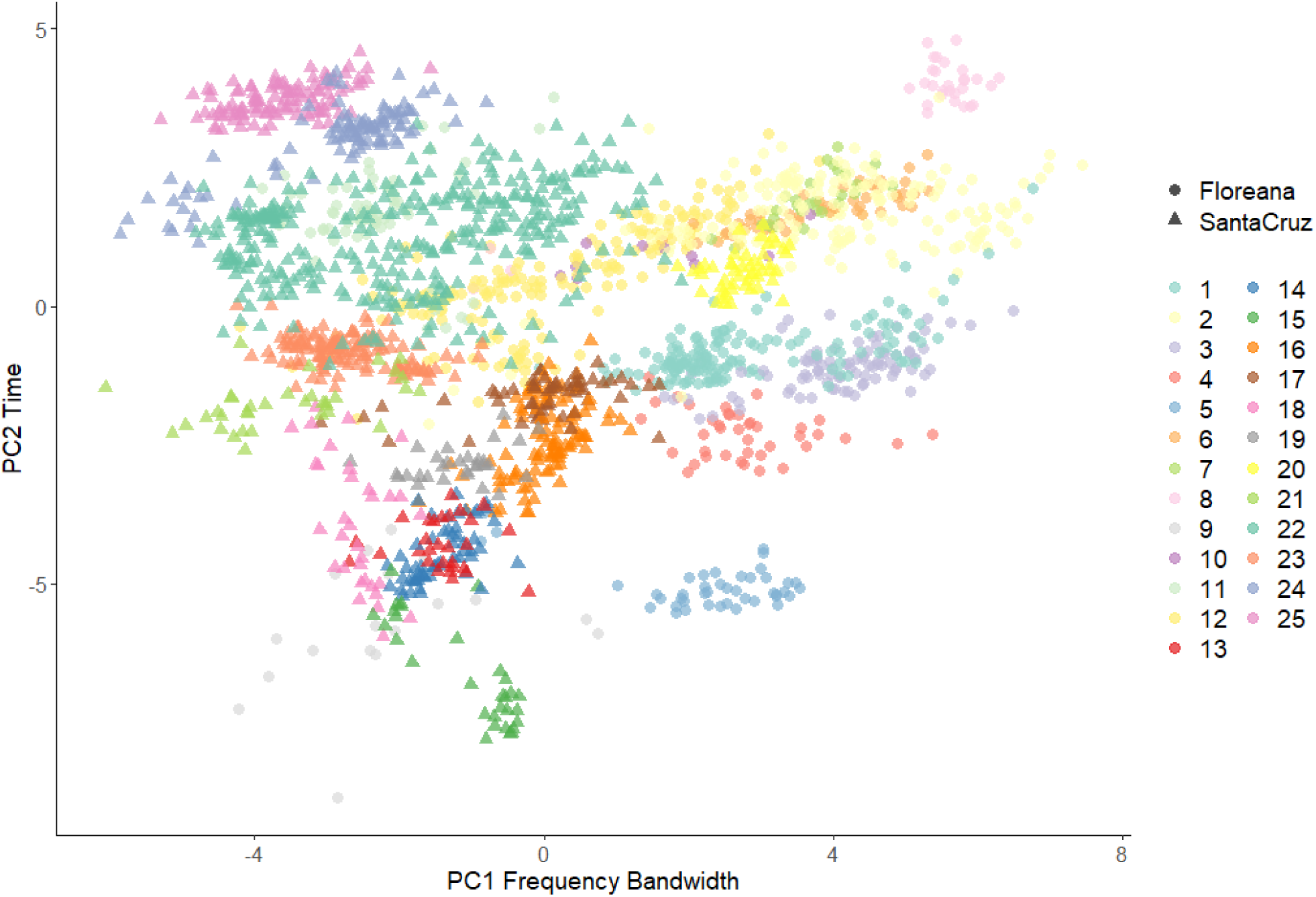
Variation in the acoustic structure of syllable types based on the first two components PC1 (dominated by frequency bandwidth) and PC2 (dominated by time-related variables) of the PCA analyzing acoustic characteristics. **A**, Santa Cruz males ● produced songs with broader frequency bandwidth than Floreana males ●. **B**, Each syllable type (1–12 for Floreana ● and 13–25 for Santa Cruz ▴) clustered in an identifiable way according to frequency bandwidth (PC1) and time (PC2).

### Differences in singing behavior

The models used to assess potential variations in male singing behavior between the islands revealed no significant differences. Males did not differ in the number of syllable types per song (estimate = 0.43, SE = 3.58, p = 0.904), the number of syllables per song (estimate = −0.004, SE = 0.09, p = 0.967), or the syllable repetition rate (estimate = −0.08, SE = 0.25, t = −0.306, p = 0.760) between the two islands.

### Differences in morphological measurements

Analysis of beak measurements (PC1_beak) showed significant differences between the islands. Cactus finch males from Santa Cruz had significantly larger beaks than males from Floreana (two-tailed t-test: t = −2.19, n = 17 of Floreana, n = 19 of Santa Cruz, p = 0.037; Table 2). There were no significant differences in body measurements (PC1_body) between the islands (two-tailed t-test: t = −1.32, n = 17 of Floreana, n = 19 of Santa Cruz, p = 0.197; Table 2).

**Table 2.** Morphological data for 17 Floreana and 19 Santa Cruz males collected between 2000 and 2024. Table shows mean values and standard error of beak size (length, depth, width in mm) and body size (tarsus, wing length in mm) per island. Beak measurements’ PCA (PC1_beak) revealed significantly larger beaks in Santa Cruz males (p = 0.037), while body size shows no significant differences between islands.

| Morphological Variables | Floreana | Santa Cruz | p |
| --- | --- | --- | --- |
| | Mean ± SE | Mean ± SE | |
| Beak length culmen | 19.5 ± 0.6 | 21.1 ± 0.4 | |
| Beak depth | 9.3 ± 0.1 | 9.6 ± 0.1 | |
| Beak width | 8.0 ± 0.2 | 8.5 ± 0.2 | |
| PC1_beak | -0.5 ± 0.4 | 0.5 ± 0.3 | **0.037** |
| Tarsus length | 22.3 ± 0.3 | 22.4 ± 0.3 | |
| Flattened wing length | 69.7 ± 0.5 | 71.2 ± 0.6 | |
| PC1_body | -0.3 ± 0.3 | 0.2 ± 0.2 | 0.197 |

## Discussion

To examine cultural and morphological divergence between two allopatric populations of Darwin’s cactus finches on Floreana and Santa Cruz Islands in the Galápagos Archipelago, we analyzed the range of song syllable types, differences in acoustic characteristics of syllables, male singing behaviors, and male morphology. Our findings support the predictions of cultural divergence in song and morphological constraints on song production, demonstrating clear differences between the two island populations. In compiling a syllable library for each population, we identified 25 syllable types across 50 males (25 from each island), with no shared syllable types between islands. Floreana males produced 12 unique syllable types, while Santa Cruz males produced 13 unique types. The absence of shared syllable types emphasizes the cultural divergence in song and supports the prediction that cultural isolation has resulted in distinct syllable repertoires in the two populations.

When populations become geographically isolated or experience bottlenecks, both genetic and cultural divergence can occur, potentially leading to song divergence within a species. The complete absence of shared syllable types between Floreana and Santa Cruz may reflect processes such as cultural innovation (the introduction of new syllables through mutations; Marler, 1970; Mundinger, 1980), cultural drift (random changes in song structure over time; (Potvin & Clegg, 2015; Williams, 2021; Williams & Lachlan, 2022), and social learning biases (Aplin, 2016) – all of which can drive rapid changes in acoustic traits (Aplin, 2019). Young Darwin’s finches acquire their songs through socially mediated learning from nearby males, typically their fathers or close neighbors (Bowman, 1979, 1983; Millington & Price, 1985; Grant & Grant, 1996, 2018; Christensen & Kleindorfer, 2007; Peters & Kleindorfer, 2018). This learning process is not error-free, similar to findings in zebra finches (Böhner, 1983, 1990; Zann, 1990; Mann & Slater, 1994). Vocal production learning involves multiple stages: initial exposure, memorization, and vocal practice leading to song crystallization (Immelmann, 1969; Marler, 1970; Marler & Peters, 1982a,b; Podos *et al*., 2009) – each susceptible to error due to perceptual inaccuracies, memory decay, or variation in motor output (Marler & Tamura, 1964; Grant & Grant, 1996; Podos & Warren, 2007). These transmission errors can accumulate and propagate within populations, leading to subtle but significant song modifications. Over generations, such changes can result in the emergence of novel syllable types or distinct population-specific song patterns (Slater, 2003; Parker *et al*., 2010; Aplin, 2016; Williams, 2021).

The cultural nature of song transmission also enables rapid adaptation to changing environmental and social contexts. Female Darwin’s finches often prefer males whose songs differ from those of their fathers, a preference that may help reduce inbreeding (Grant, 1984; Grant & Grant, 1989, 1995, 1996; Gibbs & Grant, 1989). This mechanism can also promote syllable diversity within populations, as males with uncommon or distinct song characteristics may gain a reproductive advantage (Colombelli-Négrel *et al*., 2023). As a result, song can be a flexible trait that evolves independently of genetic changes, highlighting the significant role of cultural evolution in the speciation and diversification of Darwin’s finches.

Our analysis also revealed significant differences in acoustic parameters between the two populations. Males from Floreana produced syllables with broader frequency bandwidths compared to males from Santa Cruz. These differences may be influenced by morphological constraints on song production, as beak size has been shown to limit the frequency range and syllable repetition rate. Birds with larger beaks are less capable of rapidly opening and closing their beaks, which affects their song characteristics (Podos, 2001; Podos *et al*., 2004; Huber & Podos, 2006). Consistent with this, males from Santa Cruz—who produced narrower frequency bandwidths—had larger beaks than their Floreana counterparts. This suggests that the narrower frequency bandwidths in Santa Cruz males may represent a compensatory adaptation to biomechanical limitations associated with their larger beaks while keeping the syllable repetition rate the same. Thus, song divergence may emerge as a byproduct of morphological differences in beak size (Podos, 2001; Badyaev *et al*., 2008). Apart from differences in beak size, males from Floreana and Santa Cruz showed no significant difference in overall body size. Also, no differences were found between the islands in the number of syllable types per song. However, 7 individuals (14%) in our sample produced songs with two syllable types, indicating some variation in song complexity within each population. Despite the observed differences in beak size and some acoustic parameters, no significant differences were detected in the number of syllables per song, or the syllable repetition rates between the two populations. This suggests that while the populations diverge in syllable repertoires and acoustic characteristics like frequency bandwidth, their overall singing behavior remains largely consistent. These findings suggest that the mechanisms driving song divergence are more closely related to syllable type and morphological factors rather than changes in singing behavior itself.

In addition to cultural evolution, hybridization with the closely related medium ground finch (*Geospiza fortis*) may also play a role in driving morphological and acoustic divergence between the cactus finch populations on Floreana and Santa Cruz. Previous studies have shown that interbreeding between cactus finches and medium ground finches is not uncommon (Grant & Grant, 1997a; Grant *et al*., 2003, 2004; Thackeray, 2024). Medium ground finches are characterized by shorter and broader beaks compared to cactus finches. Hybrid offspring could exhibit altered beak morphology and modified song characteristics, potentially introducing additional variability into the populations (Peters & Kleindorfer, 2018; Kleindorfer et al., 2019). This hybridization could contribute to both morphological and acoustic divergence, further complicating the mechanisms underlying song variation in the cactus finch populations.

While hybridization between cactus finches and medium ground finches likely occurs on both islands, its effects may be more pronounced in smaller populations, such as the one on Floreana (Grant *et al*., 2003). Small populations are particularly vulnerable to hybridization with closely related species due to limited mate choice and smaller gene pools (Grant & Grant, 1997a). Introgressive hybridization, as documented in Darwin’s finches, can have profound effects on morphology and behavior, including changes in beak shape and song characteristics (Grant *et al*., 2004; Kleindorfer *et al*., 2014; Peters & Kleindorfer, 2018; Dudaniec *et al*., 2025). For the cactus finches on Floreana, the introduction of medium ground finch genes could accelerate divergence, leading to distinct differences in both morphological traits and acoustic parameters when compared to larger or more stable populations, such as those on Santa Cruz.

During data collection, we also observed a potential shift in nest site preference on Floreana. Nest monitoring revealed that although *Opuntia* cacti—typically the primary nesting habitat of the cactus finch—were present, only 3 out of 25 cactus finch nests were found in *Opuntia*. Instead, most nests (15) were located in *Candelabra* cacti, while 7 were built in trees (e.g., *Palo Santo*, *Acacia*) or vines. In contrast, medium ground finches were frequently observed defending nests in *Opuntia* on Floreana, indicating possible competition or a shift in nesting preferences. On Santa Cruz, cactus finches were primarily observed defending *Opuntia*; however, comprehensive nest monitoring was not conducted on that island. The extent of this possible shift in nest site preference, as well as the genetic differences across populations, will be investigated in future research.

## Conclusion

Our findings reveal clear differences in syllable repertoires, some acoustic characteristics, and beak morphology between the two cactus finch populations, supporting the idea that both cultural and morphological factors could have shaped song divergence in this system. The absence of shared syllable types underscores the role of cultural evolution in song differentiation, while beak size constraints likely contribute to differences in syllable structure. Additionally, potential hybridization with medium ground finches may introduce further variation in song and morphology, particularly in the smaller Floreana population. Planned follow-up studies, incorporating genetic analyses and playback experiments will provide deeper insights into the mechanisms driving these differences. Such research could not only enhance our understanding of evolutionary processes and speciation but also inform conservation strategies to protect the unique ecosystems of the Galápagos Islands.

## Acknowledgements

We thank the Galápagos National Park Directorate (GNPD) for granting permission to conduct this study (PC-021-99, PC-19-07, PC-39-09, PC-58-11, PC-38-12, PC-15-14, PC-23-16, PC-02-20, PC-73-21, PC-89-22, PC-87-23; Contrato Marco MAE-DNB-CM-2016-0043), and staff at the Charles Darwin Research Station (CDRS), especially Marta Romoleroux, for providing logistical support. We thank Floreana’s GNPD office for their support, especially Eddie Rosero and his team of park rangers on Floreana Island. We thank all team members who contributed during the various field seasons, especially Bek Christensen, Lauren Common, Rachael Dudaniec, Jody O’Connor, Santiago Torres, and Carlos Vinueza, for their invaluable assistance in collecting measurements in the field. We are also grateful to the community of Floreana for their warmth and hospitality. This publication is contribution number 2743 of the Charles Darwin Foundation for the Galápagos Islands.

## Author Contributions

MK, JGL, AY, CA and SK designed the research; MK wrote the first draft of the paper; MK and JGL collected the data; MK and AY analysed the data; CA and AY provided bioacoustical advice; SK secured the funding and supervised the project; all authors edited the manuscript.

## Conflict of interest

The authors declare no conflict of interest.

## Funding

This research was funded in whole or in part by the University of Vienna’s Short-term grants abroad (KWA) to MK and by the Austrian Science Fund (FWF) [10.55776/PAT1115224], [10.55776/P36342] and [10.55776/W1262], and the Galapagos Conservation Trust Fund, with awards to SK.

## Data availability

The data for this manuscript will be made available on PHAIDRA upon acceptance.

## Supporting Information

**Table S1.**
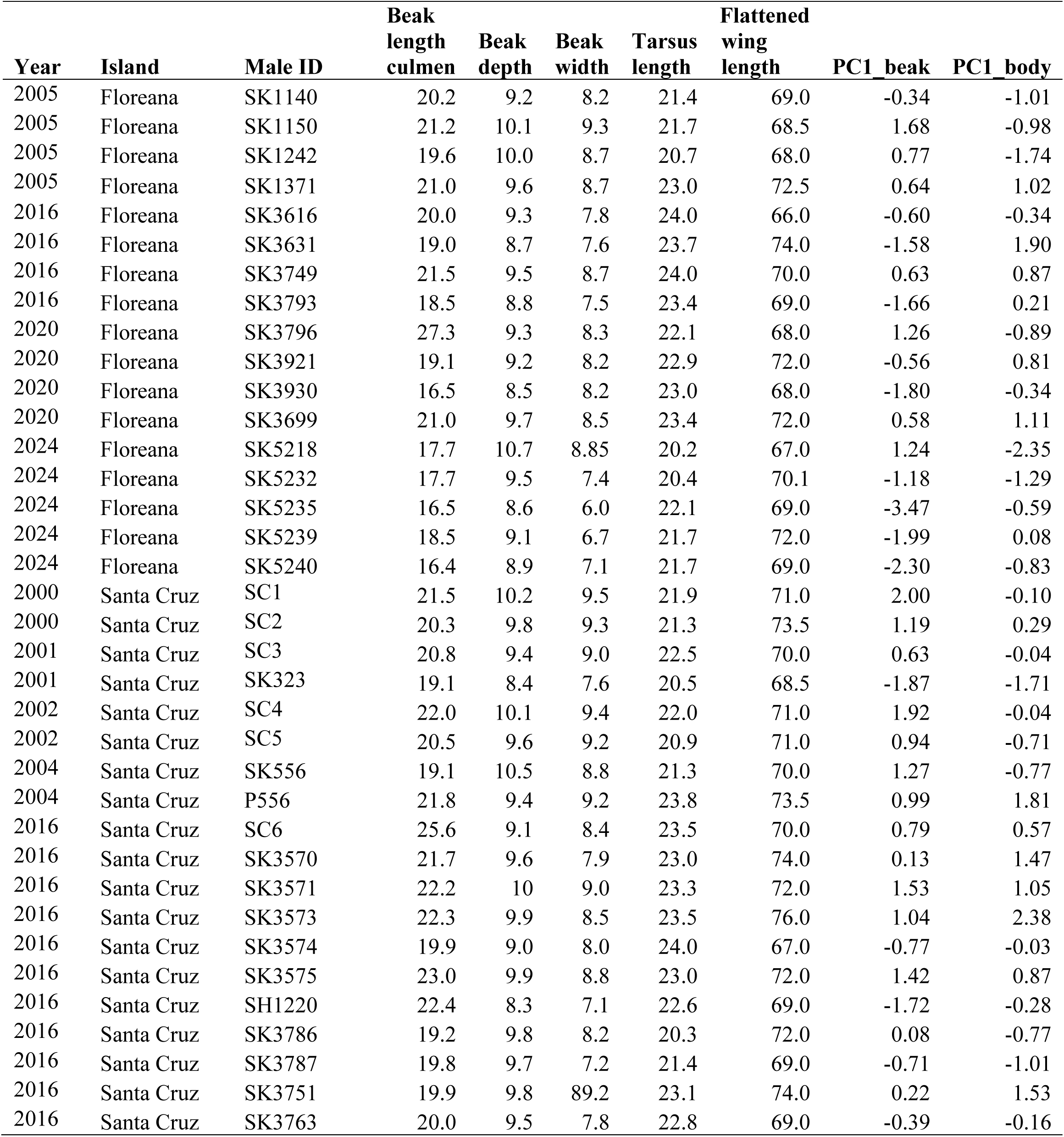
Morphological measurements of 36 male cactus finches from Floreana and Santa Cruz collected between 2000 and 2024. PC1_beak represents the first principal component from beak measurements (length, depth, width in mm), while PC1_body reflects the first principal component from body measurements (tarsus and flattened wing length in mm).

**Table S2.** Eigenvalues, proportion and cumulative variances for the first 4 components of the principal component analysis of the frequency- and time-related measurements.

| PC | Eigenvalue | Proportion<br>variance | Cumulative<br>variance |
| --- | --- | --- | --- |
| PC1 | 8.61 | 0.41 | 0.41 |
| PC2 | 6.08 | 0.29 | 0.70 |
| PC3 | 2.20 | 0.10 | 0.80 |
| PC4 | 1.18 | 0.06 | 0.86 |

**Table S3.** Factor loadings for the first 4 principal components from the PCA of 21 acoustic variables. Shading highlights the highest loadings for each PC (values > 0.3).

| Factor | PC1_freq_bandwidth<br>h | PC2_time<br>e | PC3_spectral_shape<br>e | PC4_peak_frequency<br>q |
| --- | --- | --- | --- | --- |
| Duration (s) | -0.08 | 0.38 | 0.13 | 0.12 |
| Minimum dominant frequency (kHz) | 0.04 | -0.25 | 0.27 | 0.04 |
| Maximum dominant frequency (kHz) | -0.31 | -0.04 | 0.11 | -0.16 |
| Bandwidth (kHz) | -0.31 | 0.04 | 0.02 | -0.17 |
| Peak frequency (kHz) | -0.08 | -0.08 | 0.29 | 0.53 |
| Slope | 0.16 | 0.15 | 0.02 | -0.36 |
| Mean frequency (kHz) | -0.29 | -0.15 | 0.21 | -0.04 |
| Standard deviation (kHz) | -0.30 | -0.02 | -0.08 | -0.31 |
| Frequency median (kHz) | -0.25 | -0.17 | 0.23 | 0.18 |
| Frequency Q25 (kHz) | -0.13 | -0.21 | 0.43 | 0.17 |
| Frequency Q75 (kHz) | -0.31 | -0.10 | 0.11 | -0.16 |
| Frequency IQR (kHz) | -0.31 | -0.03 | -0.05 | -0.25 |
| Time median (s) | -0.08 | 0.37 | 0.13 | 0.04 |
| Time Q25 (s) | -0.10 | 0.35 | 0.09 | 0.05 |
| Time Q75 (s) | -0.09 | 0.38 | 0.13 | 0.08 |
| Time IQR (s) | -0.08 | 0.37 | 0.14 | 0.10 |
| Skew | 0.22 | 0.02 | 0.41 | -0.33 |
| Kurtosis | 0.16 | 0.02 | 0.45 | -0.39 |
| Spectral entropy | -0.31 | 0.05 | -0.17 | 0 |
| Time entropy | 0.16 | -0.32 | 0 | -0.02 |
| Spectrographic entropy | -0.29 | -0.12 | -0.2 | -0.02 |

**Table S4.**
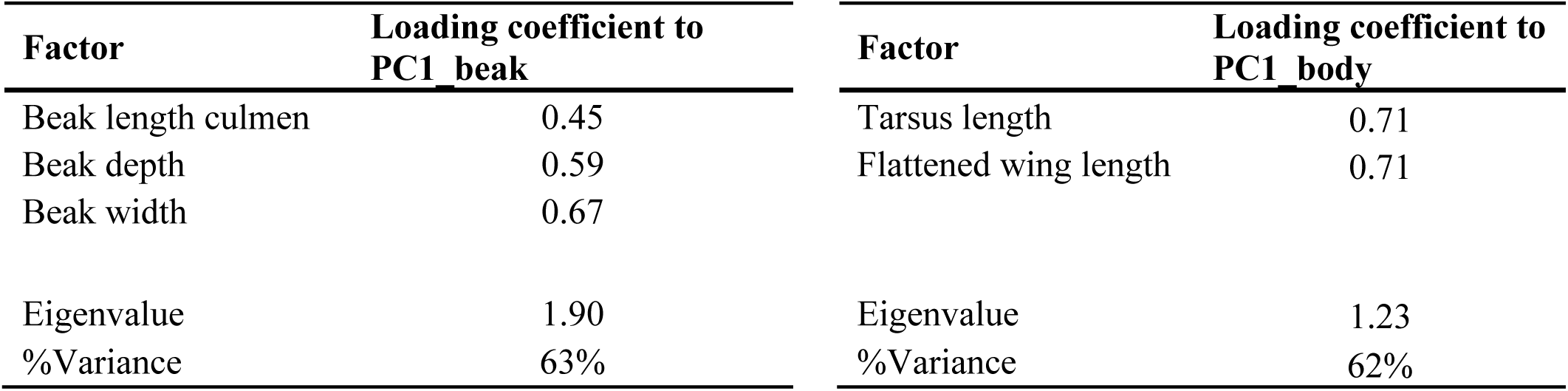
Loading coefficients of the principal component analysis of the beak and body measurements.

| Factor | Loading coefficient to<br>PC1_beak | Factor | Loading coefficient to<br>PC1_body |
| --- | --- | --- | --- |
| Beak length culmen | 0.45 | Tarsus length | 0.71 |
| Beak depth | 0.59 | Flattened wing length | 0.71 |
| Beak width | 0.67 |  |  |
| Eigenvalue | 1.90 | Eigenvalue | 1.23 |
| %Variance | 63% | %Variance | 62% |

**Table S5.** Details of seven Darwin’s cactus finch males (of 50 males recorded during 2024) that produced a song with two syllable types; all other males (86 %) had song with one syllable type. The table includes male ID and Island, the number of song recordings (n) per male, the percentage of songs with one syllable type versus two syllable types, the syllable type combinations, and three syntax examples for each male.

| Male ID | Island | #Recordings (n) | % Song with<br>1 syllable type | % Song with<br>2 syllable<br>types | Syllable types<br>present | Syntax examples |
| --- | --- | --- | --- | --- | --- | --- |
| Male 1 | Floreana | 17 | 88 | 12 | 6 & 11 | 6-6-6-6; 11-11-11; 6-6-11-11-6-6 |
| Male 2 | Floreana | 14 | 29 | 71 | 7 & 11 | 7-7; 7-11-11-11-11; 7-7-11-11-11-11-7-7 |
| Male 3 | Floreana | 10 | 0 | 100 | 10 & 8 | 10-8-8-8; 10-8-8-8-8; 10-8 |
| Male 4 | Floreana | 9 | 0 | 100 | 11 & 2 | 11-2-2-2-2-2-2; 11-2-2-2-2-11-2; 11-2-2-2-2 |
| Male 5 | Floreana | 12 | 0 | 100 | 11 & 2 | 11-2-2-2; 11-2-2; 11-2-2-2-2-2 |
| Male 6 | Santa Cruz | 23 | 78 | 22 | 24 & 13 | 24-24-24; 13-13; 24-24-24-24-24-13 |
| Male 7 | Santa Cruz | 20 | 50 | 50 | 14 & 15 | 14; 14-14-15; 14-15 |

---

**Figure S1.**
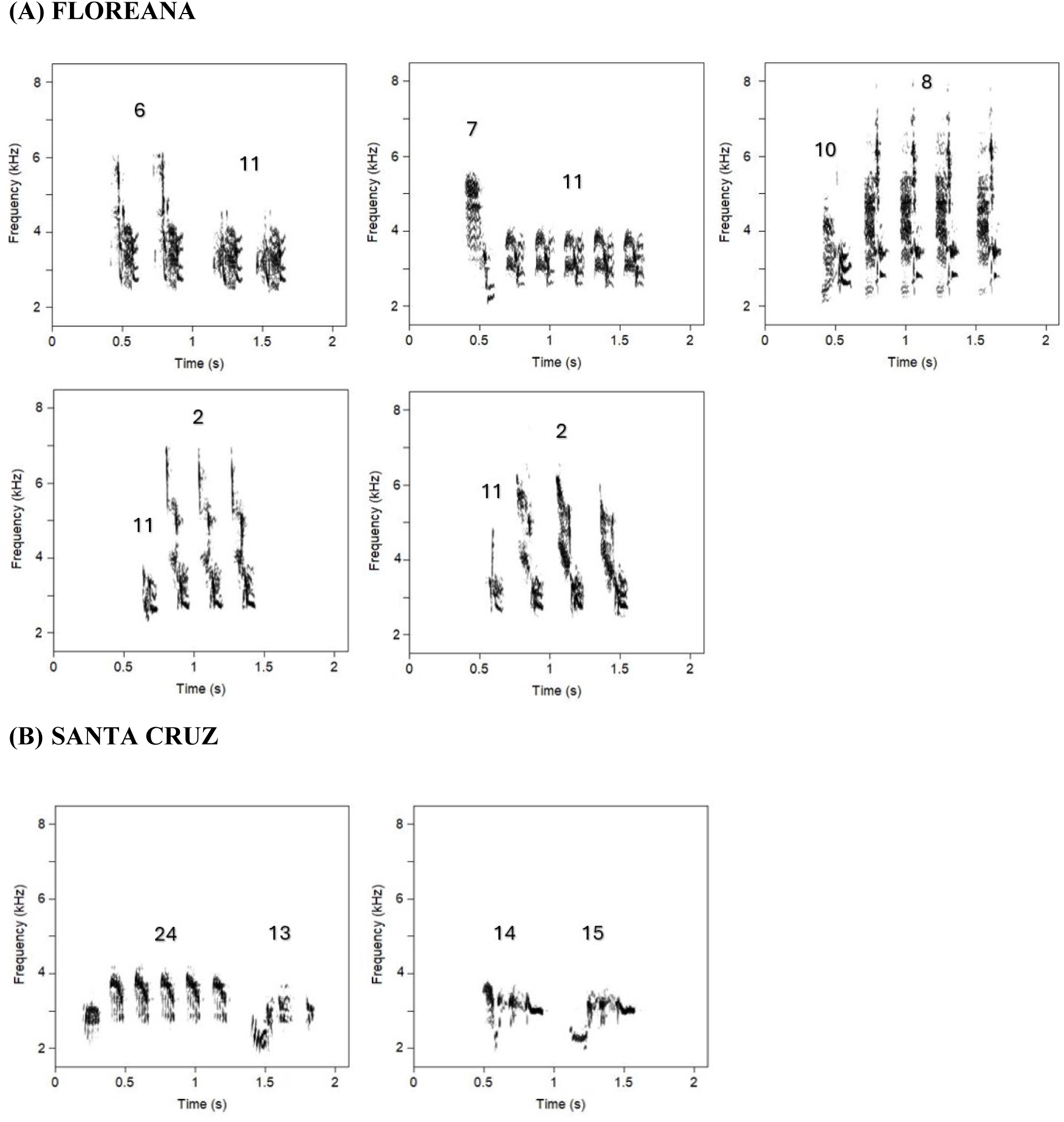
Spectrograms showcasing songs with two syllable types, including examples from the five males on **(A) Floreana** and the two males on **(B) Santa Cruz** that produced song with two syllable types in this study.

## Notes

### Competing Interest Statement

The authors have declared no competing interest.

